# In vitro rapid inactivation of SARS-CoV-2 by visible light photocatalysis using boron-doped bismuth oxybromide

**DOI:** 10.1101/2021.03.09.434359

**Authors:** Li Ling, Tea Carletti, Zihang Cheng, Ruixuan Wang, Yanxiao Ren, Paul Westerhoff, Chii Shang, Alessandro Marcello, Shuyu Chen

## Abstract

Inactivation of SARS-CoV-2 in wastewater and on surfaces is critical to prevent the fecal-oral and fomite transmission, respectively. We hypothesized that visible light active photocatalysts could dramatically enhance the rate or extent of virus inactivation and enable the use of visible light rather than shorter wavelength ultraviolet light. A novel visible light active photocatalyst, boron-doped bismuth oxybromide (B-BiOBr), was synthesized and tested for its SARS-CoV-2 inactivation towards Vero E6 cell lines in dark and under irradiation at 426 nm by a light emitting diode (LED) in water. SARS-CoV-2 inactivation in the presence of B-BiOBr (0.8 g/L) under LED irradiation reached 5.32-log in 5 min, which was 400 to 10,000 times higher than those achieved with conventional photocatalysts of tungsten or titanium oxide nanomaterials, respectively. Even without LED irradiation, B-BiOBr inactivated 3.32-log of SARS-CoV-2 in the dark due to the ability of bismuth ions interfering with the SARS-CoV-2 helicase function. LED irradiation at 426 nm alone, without the photocatalyst, contributed to 10% of the observed inactivation and was attributed to production of reactive oxygen species due to blue-light photoexcitation of molecules in the culture media, which opens further modes of action to engineer disinfection strategies. The visible light active B-BiOBr photocatalyst, with its rapid SARS-CoV-2 inactivation in the presence and absence of light, holds tremendous opportunities to build a healthy environment by preventing the fecal-oral and fomite transmission of emerging pathogens.

## Introduction

Fecal-oral transmission of the novel severe acute respiratory syndrome coronavirus 2 (SARS-CoV-2) plays a backseat to aerosol transmission, but cannot be overlooked as potential threats to the community (1, 2), because SARS-CoV-2 was found in environmental samples and can survive in wastewater over several days (3, 4). Mitigating transmission of SARS-CoV-2 in water is thus critical in both developed or developing countries. Current chemical and germicidal ultraviolet light (i.e., UV_254_) disinfection strategies can be well managed in centralized water and wastewater treatment plants to inactivate SARS-CoV-2 (3, 5). These strategies are almost impossible to be implemented at the household or community levels, where raw sewage, unsealed sewer vent pipes, leaking septic systems, medical biological wastes, etc. can all lead to SARS-CoV-2 infection (6). In some countries, mandatory disinfection, for example, chlorine-based bleach (1:99 dilution of 3% hypochlorite/hypochlorous acid in water), will be implemented upon request by the government. However, its effect is temporary, requires transport/storage of toxic chemicals, and its application is heavily labor-intensive.

Heterogeneous visible light photocatalysis (VLP) is a promising disinfection approach to rapidly inactivate virus in water. In VLP, reactive hole-electron 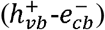 pairs are generated upon receiving visible light irradiation with light energy levels equal to or higher than bandgaps of photocatalysts (7). The 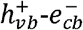 pairs then react with water and dissolved oxygen to produce reactive oxidative species (ROS), for example, hydroxyl and superoxide radicals (7). VLP were found to effectively inactivate different viruses, for example poliovirus I, hepatitis B virus, murine norovirus, and influenza virus H1N1 (8). Once coated on inner surfaces in photocatalytic reactors, VLP offers a continuous and inexpensive mode of disinfection in water without additional chemical disinfectants. While titanium dioxide is inexpensive, they require ultraviolet light (wavelengths <380 nm) to be activated (7). Even with doping to make them visible-light active (7), their viral inactivation is limited to a single mode by photocatalytic-induced ROS. Among different groups of photocatalysts, of particular interest is the bismuth-based compounds, which hold promise to be visible-light active (9) and provide a secondary inactivation mode because of the unique capability of bismuth to interfere with enzymatic functions of different microbes (10, 11).

In this study, a novel visible-light active boron-doped bismuth oxybromide (B-BiOBr) photocatalyst, with a bandgap of 2.53 eV (Fig. 1a), was synthesized and investigated for its inactivation of SARS-CoV-2 infectivity towards Vero E6 cell lines in the dark and under 426 nm LED irradiation in water (spectrum shown in Fig. 1b). The contributions of SARS-CoV-2 inactivation by B-BiOBr in the dark, under LED irradiation alone, and by photocatalytic-induced ROS towards the overall SARS-CoV-2 inactivation were revealed. The inactivation performance of the B-BiOBr photocatalysis was compared with the virus inactivation by a widely reported titanium dioxide (anatase/rutile, P25) and a tungsten oxide (WO_x_) nanocomposite photocatalysts (bandgaps shown in Fig. 1a) under 361 and 398 nm LED irradiation (spectra shown in Fig. 1b), respectively. The work also used irradiation energy normalized SARS-CoV-2 inactivation rates at different LED wavelengths to understand how the energy of different photons contribute towards the virus inactivation.

**Figure 1.**
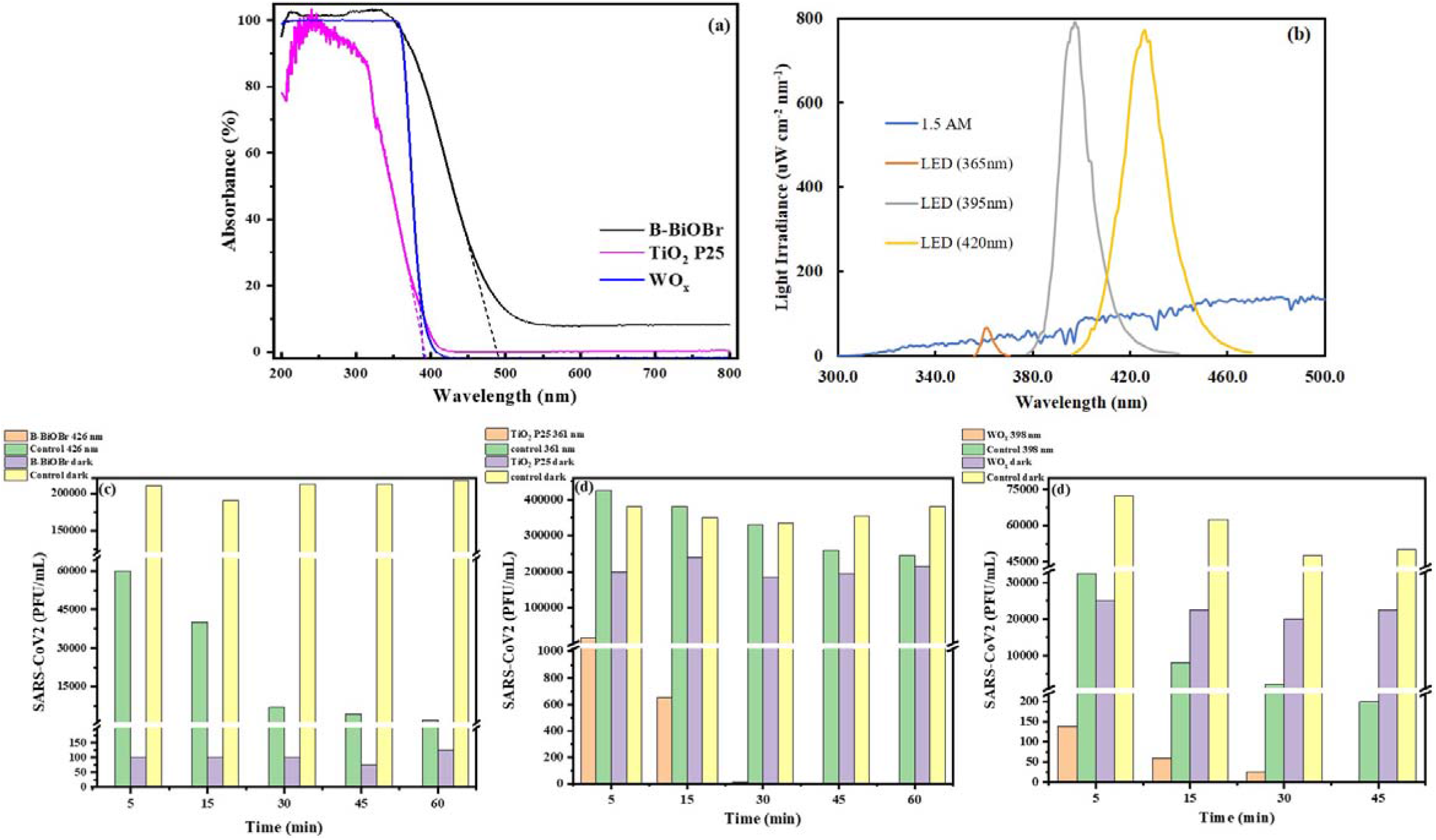
(a) UV-vis diffuse reflectance spectra of the three photocatalysts; (b) the irradiation spectra of AM 1.5, 361, 398 and 426 nm LED; (c−e) the inactivation kinetics of SARS-CoV-2 by different photocatalytic processes, condition: [B-BiOBr] = 0.8 g/L and [P25] = [WO_x_] = 5.0 g/L.

### Results and discussion

Fig. 1c, 1d and 1e shows the inactivation kinetics of SARS-CoV-2 in water by B-BiOBr, P25 and WO_x_ photocatalysis, respectively. Each of the three photocatalysis is attributable to three different processes: i) photocatalyst-mediated inactivation even without light (i.e., inactivation in the dark), ii) LED irradiation alone, and iii) photocatalytic-induced ROS inactivation. The SARS-CoV-2 inactivation is reported as log-inactivation (log inactivation = −log (N/N_0_) using the fraction remaining of infectious virus (N/N_0_)). Figure 1c shows the overall SARS-CoV-2 inactivation achieved 5.32-log with B-BiOBr (0.8 g/L) under 426 nm LED irradiation, while inactivation in the dark or by LED irradiation alone reached 3.32- and 0.54-log, respectively. The photocatalytic-induced ROS inactivation thus contributes to 1.46-log, as calculated by subtracting the overall SARS-CoV-2 inactivation with those in the dark and by LED irradiation. The contributions of three individual processes to the photocatalytic SARS-CoV-2 inactivation by B-BiOBr follow: inactivation in the dark (62%) > photocatalytic-induced ROS inactivation (28%) > LED irradiation alone (10%). A significant percentage of the SARS-CoV-2 inactivation by the B-BiOBr photocatalysis occurred in the dark. Surface adsorption was unlikely the major reason, because the BET surface area of B-BiOBr (3.4 m^2^/g) is an order of magnitude lower than for P25 (55.5 m^2^/g) (SI), where the later catalysts showed little virus inactivation in the dark (Fig. 1d). We hypothesized the much stronger inactivation of SARS-CoV-2 by B-BiOBr in the dark shares a similar mechanism proposed by Yuan et al. (2020): Ranitidine bismuth citrate suppressed SARS-CoV-2 replication and relieves virus-associated pneumonia in Syrian hamsters by replacing Zn(II) on the SARS-CoV-2 helicase (Nsp13) with bismuth ions (11). We also found that not all photocatalysts would be as potent as BiOBr. The overall 5-min SARS-CoV-2 inactivation by P25 under 361 nm (5 g/L, Fig. 1d) and WO_x_ under 398 nm (5 g/L, Fig. 1e) were just 1.32-log and 2.71-log, respectively. For WO_x_ and P25 to achieve 4- to 5-log SARS-CoV-2 inactivation, respectively, requires much longer time (45 min) with the same power input (5 W) of different LEDs.

In the absence of any photocatalyst, SARS-CoV-2 could be inactivated using 361, 398 and 426 nm light emitted from different LEDs. Inactivation could be fitted with pseudo first-order kinetics with rate constants of 5.2 × 10^−5^, 86 × 10^−5^ and 63 × 10^−5^ s^−1^ using 361, 398 and 426 nm LEDs, respectively. The corresponding irradiation energy normalized inactivation rate constants were 0.045, 0.019 and 0.0097 µE^−1^ cm^2^, with higher values at shorter wavelength. A prior work attributed the inactivation of Feline calicivirus irradiated at 405 nm light to the generation of ROS by blue-light photosensitizers in the culture media, not to absorption of photons by the virus (12). The presence of UV-A/blue-light photosensitizers in saliva and the aqueous environment (13) opens up further modes of action to engineered light sources using UV-A/blue-light instead of UV_254_ light, which damages human skin and eyes when directly viewed, and oxidizes plastics or other surfaces with prolonged exposures (14). But the feasibility of such practice needs further investigation. We also simulated that 0.9-log inactivation of SARS-CoV-2 is expected in 30 min by solar irradiation reaching the earth’s surface (AM 1.5 standard spectrum), which shows a stronger inactivation than that in artificial saliva in the dark (about 0.3-log in 30 min) (15), but is still insufficient in many realistic scenarios.

Our current results illustrate the excellent performance of the B-BiOBr photocatalyst, whose SARS-CoV-2 inactivation rate is of practical significance to prevent fecal-oral transmission. The unique capability of bismuth interfering with microbial enzymatic functions provides B-BiOBr a secondary dark inactivation mode, in addition to the conventional visible-light photocatalytic-induced ROS inactivation mode, towards different microbes. Once coated on frequently touched surfaces, the B-BiOBr photocatalysis has the potential to serves as a preventive or an alternative strategy to prevent hand-to-mouth fomite transmission in public settings, where continuous application of chlorine or UV_254_ light disinfection strategies are not recommended or impossible. The dual modes of action for visible light active B-BiOBr photocatalyst holds tremendous opportunities to build a healthy environment by preventing the fecal-oral and potentially fomite transmission of emerging pathogens.

### Materials and methods

B-BiOBr was synthesized via a hydrothermal method using boric acid, nitric acid, bismuth nitrate tetrahydrate, cetyltrimethylammonium bromide, and sodium hydroxide. The elemental mass % of B-BiOBr was B:Bi:O:Br:C = 0.51:73.53:5.32:17.15:3.49 as identified using XPS (SI). WO_x_ was synthesized following a previous publication (16). P25 was purchased from Sigma-Aldrich. All chemicals were used without further treatment. The experimental setup used for the SARS-CoV-2 inactivation and the detailed testing procedures are shown in SI. Detailed calculations of the fluence normalized inactivation rates and the SARS-CoV-2 inactivation rates under AM 1.5 standard spectrum are shown in SI.

SI information will be provided upon request.

## Acknowledgement

This work was partially funded by Raze Technology limited, the Hong Kong Research Grants Council (T21-604/19-R), and the National Science Foundation (EEC-1449500) Nanosystems Engineering Research Center on Nanotechnology-Enabled Water Treatment. Some of the authors are coinventors on a US provisional patent with application no. 63158378. Conflicts of interest are managed by Dr. Shuyu Chen.

## Notes

### Competing Interest Statement

This work was partially funded by Raze Technology Ltd., the Hong Kong Research Grants Council (T21-604/19-R), and the National Science Foundation (EEC-1449500) Nanosystems Engineering Research Center on Nanotechnology-Enabled Water Treatment. Some of the authors are coinventors on a US provisional patent with application no. 63158378. Conflicts of interest are managed by Dr. Shuyu Chen. Shuyu Chen is the board of director in Raze Technology Company.

## References

1. P.Y. Chia, K.K. Coleman, Y.K. Tan, S.W.X. Ong, M. Gum, S.K. Lau, X.F. Lim, A.S. Lim, S. Sutjipto, P.H. Lee, T.T. Son, B.E. Young, D.K. Milton, G.C. Gray, S. Schuster, T. Barkham, P.P. De, S. Vasoo, M. Chan, B.S.P. Ang, B.H. Tan, Y.S. Leo, O.T. Ng, M.S.Y. Wong, K. Marimuthu, D.C. Lye, P.L. Lim, C.C. Lee, L.M. Ling, L. Lee, T.H. Lee, C.S. Wong, S. Sadarangani, R.J. Lin, D.H.L. Ng, M. Sadasiv, T.W. Yeo, C.Y. Choy, G.S.E. Tan, F. Dimatatac, I.F. Santos, C.J. Go, Y.K. Chan, J.Y. Tay, J.Y.L. Tan, N. Pandit, B.C.H. Ho, S. Mendis, Y.Y.C. Chen, M.Y. Abdad, D. Moses, Detection of air and surface contamination by SARS-CoV-2 in hospital rooms of infected patients. Nat. Commun. 11, 1–7 (2020).

2. World Health Organization, Transmission of SARS-CoV-2: implications for infection prevention precautions. 2020. https://www.who.int/news-room/commentaries/detail/transmission-of-sars-cov-2-implications-for-infection-prevention-precautions

3. A.W.H. Chin, J.T.S. Chu, M.R.A. Perera, K.P.Y. Hui, H.-L. Yen, M.C.W. Chan, M. Peiris, L.L.M. Poon, Stability of SARS-CoV-2 in different environmental conditions. The Lancet Microbe 1 (1), E10 (2020).

4. A. Bivins, J. Greaves, R. Fischer, K.C. Yinda, W. Ahmed, M. Kitajima, V.J. Munster, K. Bibby, Persistence of SARS-CoV-2 in water and wastewater. Environ. Sci. Technol. Lett. 7 (12), 937–942 (2020).

5. C.P. Sabino, F.P. Sellera, D.F. Sales-Medina, R.R.G. Machado, E.L. Durigon, L.H. Freitas-Junior, M.S. Ribeiro, UV-C (254 nm) lethal doses for SARS-CoV-2. Photodiagn. Photodyn. Ther. 32, 101995 (2020).

6. M. Usman, M. Farooq, K. Hanna, Existence of SARS-CoV-2 in wastewater: Implications for its environmental transmission in developing communities. Environ. Sci. Technol. 54 (13), 7758–7759 (2020).

7. S.K. Loeb, P.J.J. Alvarez, J.A. Brame, E.L. Cates, W. Choi, J. Crittenden, D.D. Dionysiou, Q. Li, G. Li-Puma, X. Quan, D.L. Sedlak, D.T. Waite, P. Westerhoff, J.-H. Kim, The technology horizon for photocatalytic water treatment: sunrise or sunset?. Environ. Sci. Technol. 53, 2937–2947 (2018).

8. C. Zhang, Y. Li, D. Shuai, Y. Shen, D. Wang, Progress and challenges in photocatalytic disinfection of waterborne viruses: A review to fill current knowledge gaps. Chem. Eng. J. 355, 399–415 (2019).

9. R. Wang, T.P. Lai, P. Gao, H. Zhang, P.L. Ho, P.C.Y. Woo, G. Ma, R.Y.T. Kao, H. Li, H. Sun. Bismuth antimicrobial drugs serve as broad-spectrum metallo-β-lactamase inhibitors. Nat. Commun. 9, 439 (2018).

10. W. Wang, G. Huang, J.C. Yu, P.K. Wong. Advances in photocatalytic disinfection of bacteria: Development of photocatalysts and mechanisms. J. Environ. Sci. (China) 34 (1), 232–247 (2015).

11. S. Yuan, R. Wang, J.F.-W. Chan, A.J. Zhang, T. Cheng, K.K.-H. Chik, Z.-W. Ye, S. Wang, A.C.-Y. Lee, L. Jin, H. Li, D.-Y. Jin, K.-Y. Yuen, H. Sun, Metallodrug ranitidine bismuth citrate suppresses SARS-CoV-2 replication and relieves virus-associated pneumonia in Syrian hamsters. Nat. Microbiol 5, 1439–1448 (2020).

12. R.M. Tomb, M. Maclean, J.E. Coia, E. Graham, M. McDonald, C.D. Atreya, S.J. MacGregor, J.G. Anderson, New proof-of-concept in viral inactivation: Virucidal efficacy of 405 nm light against feline calicivirus as a model for norovirus decontamination. Food Environ Virol. 9 (2), 159–167 (2017).

13. D. Zhang, S. Yan, W. Song, Photochemically induced formation of reactive oxygen species (ROS) from effluent organic matter. Environ. Sci. Technol. 48 (21), 12645–12653 (2014).

14. F.R. De Gruijl, H.J.C.M. Sterenborg, P.D. Forbes, R.E. Davies, C. Cole, G. Kelfkens, H. Van Weelden, H. Slaper, J.C. Van Der Leun, Wavelength dependence of skin cancer induction by ultraviolet irradiation of albino hairless mice. Cancer Res. 53 (1), 53–60 (1993).

15. S.J. Smither, L.S. Eastaugh, J.S. Findlay, M.S. Lever, Experimental aerosol survival of SARS-CoV-2 in artificial saliva and tissue culture media at medium and high humidity. Emerg. Microbes Infect. 9 (1), 1415–1417 (2020).

16. C. Wang, A. Li, C. Li, S. Zhang, H. Li, X. Zhou, L. Hu, Y. Feng, K. Wang, Z. Zhu, R. Shao, Y. Chen, P. Gao, S. Mao, J. Huang, Z. Zhang, X. Han, Ultrahigh photocatalytic rate at a single[metal[atom[oxide. Adv. Mater. 31, 1903491 (2019).

