## Supplementary material for "In vitro rapid inactivation of SARS-CoV-2 by visible light photocatalysis using boron-doped bismuth oxybromide": SI

**Submitted to:**

Proceedings of the National Academy of Sciences of the United States of America

Li Ling^1,*^, Tea Carletti^2^, Zihang Cheng^1^, Ruixuan Wang^1^, Yanxiao Ren^3^, Paul Westerhoff^4^, Chii Shang^1,5^, Alessandro Marcello^2^, Shuyu Chen^6,*^

1. Department of Civil and Environmental Engineering, the Hong Kong University of Science and Technology, Clear Water Bay, Kowloon, Hong Kong

2. Laboratory of Molecular Virology, International Centre for Genetic Engineering and Biotechnology (ICGEB), Trieste, Italy

3. Department of Physics, the Hong Kong University of Science and Technology, Clear Water Bay, Kowloon, Hong Kong

4. Nanosystems Engineering Research Center for Nanotechnology-Enabled Water Treatment, School of Sustainable Engineering and The Built Environment, Arizona State University, Tempe, AZ, United States

5. Hong Kong Branch of Chinese National Engineering Research Center for Control & Treatment of Heavy Metal Pollution, the Hong Kong University of Science and Technology, Clear Water Bay, Kowloon, Hong Kong

6. Raze Technology limited, Genesis, 33-35 Wong Chuk Hang Road, Hong Kong

**Extended materials and methods:**

**Brunauer–Emmett–Teller (BET) surface area analysis of B-BiOBr and TiO_2_:**

BET surface area analysis was conducted using JW-BK200C. 0.7433 g of B-BiOBr and 0.0738 g of TiO_2_ were pretreated at 110 degree Celsius for 10 h. The adsorption temperature was 77 K. The multi-point BET surface area result of B-BiOBr and TiO_2_ were 3.39 and 55.5 m^2^/g, respectively.

**X-ray photoelectron spectroscopy results of B-BiOBr:**

Three samples of B-BiOBr were analyzed by an X-ray photoelectron spectroscopy (Physical Electronics 5600). The results are summarized in the table below. An average of those data was taken to obtain the elemental mass composition of B-BiOBr.

|  | Sample 1 (%) | Sample 2 (%) | Sample 3 (%) | Average (%) |
| --- | --- | --- | --- | --- |
| O 1s | 5.77 | 5.34 | 4.84 | 5.32 |
| C 1s | 4.90 | 2.93 | 2.65 | 3.49 |
| B 1s | 0.40 | 0.56 | 0.56 | 0.51 |
| Bi 4f | 74.53 | 73.23 | 72.83 | 73.53 |
| Br 3d | 14.40 | 17.94 | 19.12 | 17.15 |

**Experimental Setup:**

An LED lamp was fixed on a stand. Four cylindrical reactors with a diameter and depth of 3.5 and 2 cm, respectively, containing 20 mL of solution, were placed 10 cm below the LED lamp on a stir plate. Three 5 W LED lamps with different light spectra were selected. The light spectra of the LED lamps are peaked at 361 nm, 398 nm and 426 nm, which are denoted as 361 nm LED, 398 nm LED, and 426 nm LED, respectively. The irradiation spectra and intensities of the three LEDs were measured by NIST Traceable Light 142 Measurement Systems (InternationalLight Technologies) at the surface of the reacting solutions.

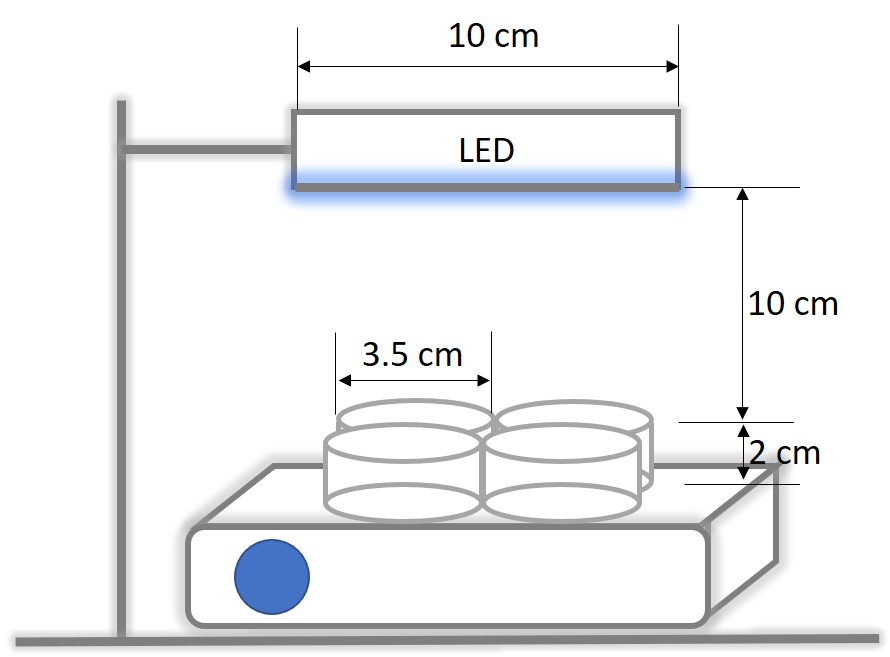

Figure S1. Schematic of experimental setup.

**Experimental Procedures:**

Vero E6 cells are used. Plastics materials used include cell culture dishes and plates, tubes, pipettes, gloves, tips, etc…Reagents for cell culture include culture medium DMEM (Dulbecco’s Modified Eagle’s Medium), foetal bovine serum (FBS), phosphate buffered saline (PBS), trypsine. SARS-CoV-2 was using isolate FVG_ICGEB_S5 reported in Licastro et al. (2020). Reagents for plaque assay include carboxymethyl cellulose 8% (CMC), paraformaldehyde 3,7% (PFA), crystal violet 1%, water. The experiments were all done in cell culture room and BSL3 facility.

Plaque Assay: Vero E6 cells were seeded in 24-well plates in order to get a full confluence the second day. Virus was serially diluted 10-fold in DMEM. 200 μl of the solution were added to each well and incubated for 1 h at 37 °C. Virus was then removed and 1 ml of overlay medium (1.5% carboxymethyl cellulose in DMEM supplemented with 2% heat inactivated FBS) was added to each well. The plates were incubated at 37 °C under 5% CO2 for 3 days. After the removal of the overlay medium, the cells were fixed with 3.7% paraformaldehyde solution for 20 min and stained with 1% crystal violet solution for 20 min followed by extensive washing with distilled water. Plaques were manually counted and multiplied by the dilution factor to determine the virus titre at the unit of plaque-forming units per milliliter (PFU/ml).

**Detailed calculations of the fluence normalized inactivation rates:**

Light irradiances of the three LEDs were measured by NIST Traceable Light Measurement Systems (InternationalLight Technologies) at the surface of the reacting solutions. The peak wavelength and half width of the 361 nm LED are 361 and 6 nm, respectively, those of the 398 nm LED are 398 and 14 nm, respectively, and those of the 426 nm LED are 426 and 20 nm, respectively. The fluence at each wavelength of the three LEDs ($f_{\lambda_{i}}$) were calculated from the following equation:

$$f_{\lambda_{i}}=\frac{I_{i}\times{10}^{-6}}{\frac{hc}{\lambda}\times1eV\times N_{A}}$$

where $I_{i}$ is the light intensity at each wavelength, h is the plank constant, c is the speed of light, λ is the wavelength in nm, 1 eV is the electron volt, and $N_{A}$ is the Avogadro constant. The fluences of the 361, 398 and 426 nm LEDs were the sum of their fluence at each wavelength. The fluence normalized inactivation rates were then calculated by dividing pseudo first order SARS-CoV-2 inactivation rates of the 361, 398 and 426 nm LEDs with their corresponding fluences at 361, 398 and 426 nm. The fluence normalized inactivation rates were thus 4.48 × 10^-2^, 1.88 × 10^-2^ and 9.73 × 10^-3^ µE^-1^ cm^2^ under 361, 398 and 426 nm light irradiation, respectively.

**Detailed calculations of the SARS-CoV-2 inactivation rate under AM 1.5 standard spectrum:**

Based on the calculated fluence normalized inactivation rates at 361, 398 and 426 nm, the SARS-CoV-2 inactivation rate under AM 1.5 standard spectrum was calculated. First, the fluence of the AM 1.5 standard spectrum at each wavelength from 356 to 470 nm were calculated following the same protocol discussed in the section of “**Detailed calculations of the fluence normalized inactivation rates**”. Then, the SARS-CoV-2 inactivation rates at 361±3, 398±7, and 426±10 nm were calculated by multiplying the fluence of the AM 1.5 standard spectrum at each wavelength range with their corresponding fluence normalized inactivation rates. The overall SARS-CoV-2 inactivation rate under AM 1.5 standard spectrum was thus the sum of the three SARS-CoV-2 inactivation rates at 361±3, 398±7, and 426±10 nm.
